# Jade1S SUMOylates tau and targets oligomers for degradation

**DOI:** 10.64898/2026.11.03.764021

**Authors:** Soong Ho Kim, Vanessa L Stahl, Lyucheng Zou, Samin Hassan, Herbert T Cohen, Kurt Farrell, Richard W Vachet, Jennifer N Rauch, John F Crary

## Abstract

The accumulation of insoluble tau aggregates is a pathological hallmark of tauopathies, including Alzheimer’s disease. While molecular chaperones and clearance pathways regulate tau homeostasis, the mechanisms governing tau oligomerization and degradation remain incompletely defined. Recent genome-wide association studies (GWAS) have linked the *JADE1* locus to tauopathy, and the PHD zinc finger protein Jade1S has been identified in complexes with tau in post-mortem brain tissue, prompting investigation into their biochemical relationship. Here, we characterize the reciprocal interactions between Jade1S and 0N4R tau stability and turnover. Covalent labeling mass spectrometry and co-immunoprecipitation mappings reveal that Jade1S binds wild-type 0N4R tau through redundant sites within its N-terminal and plant homeodomain (PHD) zinc finger domains. *In vitro* assays demonstrate that Jade1S functions as a SUMO E3 ligase, utilizing its PHD domains to mediate tau SUMOylation. Cell-based cycloheximide chase assays reveal a reciprocal stabilization dynamic: tau co-expression extends the overall physiological half-life of Jade1S from less than an hour to ∼17 hours, arresting its turnover specifically within sarkosyl-insoluble fractions. Conversely, Jade1S selectively accelerates the degradation of sarkosyl-insoluble tau without altering the turnover of the soluble monomeric pool over chased time. Mechanistically, Jade1S redirects the clearance of SUMO2/3-modified tau oligomers away from basal autophagic-lysosomal pathways and toward proteasome-dependent degradation. Furthermore, Jade1S exhibits intrinsic molecular chaperone activity, inhibiting arachidonic acid-induced tau oligomerization *in vitro*. Together, these data characterize Jade1S as a dual-function SUMO ligase and chaperone that regulates tau stability, prevents oligomer assembly, and promotes the targeted proteasomal clearance of aggregated tau.

## Introduction

In healthy neurons, tau exists in a dynamic equilibrium between a microtubule-bound state and a cytosolic pool (Konzack, Thies et al. 2007, Breuzard, Hubert et al. 2013). Post-translational modifications and pathogenic mutations drive tau detachment from microtubules, significantly increasing free-floating cytosolic tau or causing it to misfold and assemble into insoluble aggregates (Hong, Zhukareva et al. 1998, Alonso, Zaidi et al. 2001, Cohen, Guo et al. 2011). The accumulation of neurofibrillary tau tangles (NFTs) is a defining hallmark of tauopathies, including Alzheimer’s disease (AD), progressive supranuclear palsy (PSP), and frontotemporal dementia (FTD). In the adult human brain, alternative splicing of the *MAPT* gene yields six major tau isoforms containing zero, one, or two N-terminal inserts (0N, 1N, 2N) and three or four microtubule-binding repeats (3R, 4R). Tauopathies are classified by the predominant isoforms within these aggregates: 3R (e.g., FTD), 4R (e.g., PSP), or 3R/4R (e.g., AD) (Creekmore, Watanabe et al. 2024).

To counter pathological misfolding and aggregation, cells maintain multiple defense mechanisms. Native monomeric tau adopts a compact "paper-clip" conformation where its N- and C-termini fold back over the central microtubule-binding domain, physically shielding aggregation-prone motifs (Jeganathan, von Bergen et al. 2006). Kinase– phosphatase homeostasis preserves tau’s positive charge to maintain microtubule binding capability (Stoothoff and Johnson 2005, Liu, Li et al. 2007, Yu, Run et al. 2009), while molecular chaperones (Hsp70, Hsp90, sHsps) monitor protein folding (Dou, Netzer et al. 2003, Karagoz, Duarte et al. 2014, Baughman, Clouser et al. 2018). Ubiquitin ligases such as CHIP clear excess or aberrantly modified tau via the ubiquitin-proteasome system prior to nucleation (Petrucelli, Dickson et al. 2004, Shimura, Schwartz et al. 2004, Dickey, Kamal et al. 2007), and enzymes such as TRIM11 and HDAC6 maintain tau solubility while actively disassembling preformed fibrils (Balmik, Sonawane et al. 2021, Zhang, Harischandra et al. 2023).

Despite this protective machinery, chronic pathological stress overrides cellular defenses, underscoring the need to identify additional regulatory factors. Jade1S (Jade Family PHD Finger 1, short isoform), a protein known for its roles in chromatin modification, cell division, and tumor suppression (Chitalia, Foy et al. 2008, Siriwardana, Meyer et al. 2015, Borgal, Rinschen et al. 2016), has recently been implicated in tauopathy pathogenesis. Genome-wide association studies (GWAS) linked *JADE1* locus variants to primary age-related tauopathy (PART). In post-mortem brain tissue from 4R and 3R/4R tauopathy cases, Jade1S localizes to neurons containing mature NFTs and physically forms a protein complex with 0N4R tau in brain lysates (Farrell, Kim et al. 2022). Single-soma RNA sequencing revealed that *JADE1* is upregulated in tangle-bearing neurons, reflecting an endogenous protective response. Supporting this hypothesis, knockdown of the *Drosophila JADE1* homolog *rhinoceros* (*rno*) significantly exacerbates tau toxicity *in vivo*. Herein, using cell-free *in vitro* assays and HEK293 cell models, we present a detailed study into how Jade1S and Tau reciprocally affect each other’s stability and turnover.

## Materials and Methods

### DEPC labeling

Each protein (20 μM) was reacted with 8-fold molar excess of DEPC at 37 °C for 1 minute. The reactions were quenched with 50-fold molar excess of imidazole to DEPC. The excess imidazole and any remaining DEPC was then removed from the samples using 10 kDa Millipore Amicon spin filters. After speed-vac drying, samples were denatured with 6 M guanidine hydrochloride, reduced with 85 mM dithiothreitol at 95 °C for 10 minutes, and alkylated with 140 mM iodoacetamide at room temperature in the dark for 30 minutes. Samples were enzymatically digested with a trypsin/Lys-C (1-1.25 μg) mixture while shaking overnight at 37 °C. The digestion was terminated with 0.5 μL of trifluoracetic acid (TFA). Samples were then cleaned up with Pierce^TM^ C-18 spin columns before LC-MS/MS analysis.

### Liquid chromatography tandem mass spectrometry (LC-MS/MS) analysis

Reversed-phase HPLC separations using an Acclaim C18 PepMap column (2 μm, 100Å, 0.3 × 150mm; Thermo Scientific) were carried out on a Dionex Ultimate 3000 system. A flow rate of 4 μL/ min was used, and a gradient of LC-MS-grade water with 0.1% formic acid (solvent A) and LC-MS-grade acetonitrile with 0.1% formic acid (solvent B) was performed using the following conditions. For the first 5 minutes, the mobile phase consisted of 95% solvent A and 5% solvent B and was directed to waste for desalting. The next 50 minutes were then coupled to a Thermo Scientific Orbitrap Fusion Tribrid for online LC-MS/MS analysis using a linear mobile phase gradient from 5% B to 50% B. The column was then washed with 95% B and re-equilibrated to 5% B over the final 20 minutes.

Mass spectra on the Orbitrap were collected in positive mode with a resolution of 60,000. The electrospray source was set to 4,000 V. Collision-induced dissociation (CID) on the quadrupole ion trap was set at 35% collision energy to acquire tandem mass spectra. Additionally, a peak inclusion list (PIL) consisting of all possible labeled and unlabeled peptides was used. Data dependent scans of the most abundant ions were also performed even if the ions were not on the PIL.

### CL-MS data analysis

A custom software pipeline for DEPC CL-MS experiments was used to identify DEPC labeling sites and to quantify the extent of labeling (Borotto, Zhou et al. 2015). Data processing search parameters were set to 10 ppm precursor mass tolerance and a 0.5 Da tolerance for product ions. Carbamidomethylation of Cys, as well as the DEPC modification of nucleophilic residues (Cys, His, Lys, Ser, Thr, Tyr) and the N-terminus were set as variable modifications. Peptides with up to three missed cleavages were considered.

Reconstructed ion chromatograms were used to calculate the extent of DEPC modification (% modification) (Eq. 1) based on the peak areas of the labeled and unlabeled ions.

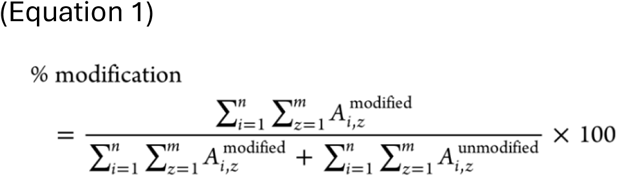

The peak area (A_i,z_) of any detected peptide (*i*) and all charge states (*z*) of that peptide for each residue of interest were considered. It is assumed that all charge states that are summed are present in all samples. Since there is a difference in ionization efficiency between unmodified and modified peptides, resulting in different retention times, the reported % modifications are relative values.

### Recombinant Jade1S and Jade1S-dd production

Competent *E. coli* BL21 cells (New England Biolabs) were transformed with pGEX-6P-1-Jade1S or pGEX-6P-1-Jade1S-dd constructs (Zhou, Wang et al. 2002, Chitalia, Foy et al. 2008). Bacteria were grown in Terrific Broth (Fisher Scientific) supplemented with 100 µg/mL carbenicillin (Fisher Scientific). Protein expression was induced in the early log phase (OD600 ∼1.0) with 100 µM isopropyl-β-D-thiogalactopyranoside (IPTG) (EMD Millipore) at room temperature for 16 h. Bacteria were lysed in a buffer containing 50 mM Tris-HCl, pH 8.0, 1% (v/v) Triton X-100, 1 mg/mL lysozyme (Sigma Aldrich), 150 mM NaCl, 1 mM EDTA, Halt protease inhibitor cocktail (Thermo Scientific), and Benzonase Nuclease (EMD Millipore). The lysate was incubated on a vertical rotator at 4 °C for 30 min, followed by incubation on ice for 30 min with brief vortexing every 5 min. The supernatant was collected following centrifugation at 16,000 × *g* for 20 min, mixed with Glutathione Sepharose 4B beads (Cytiva), and incubated at 4 °C overnight. Jade1S or Jade1S-dd was cleaved from the bead-bound GST tags using PreScission protease (Cytiva/GenScript) according to the manufacturer’s protocol.

### In vitro SUMOylation assay

SUMOylation reactions were reconstituted in a 20 µL volume using ubiquitination buffer (20 mM Tris-HCl, pH 7.6, 50 mM NaCl, 1 mM β-mercaptoethanol, and 5 mM MgCl2; UBPBio). The reactions contained 50 nM 6xHis-SAE1/SAE2 (UBPBio), 200 nM 6xHis-UbE2I (UBPBio), 17.2 μM SUMO1 (UBPBio) or 36.7 μM SUMO2 (UBPBio), 100 nM 0N4R tau (rPeptide), 2 mM ATP (UBPBio), 5% (v/v) glycerol, and Jade1S or Jade1S-dd (0, 125, 250, 500, or 1000 nM). Reactions were incubated at 37 °C for 3 h.

### Cell culture and transfections

HEK293 cells were obtained from ATCC (CRL-1573) and cultured in Dulbecco’s Modified Eagle Medium (DMEM) (Gibco 11965-092) supplemented with 10% fetal bovine serum (FBS) (Gibco A5670801) and 1% penicillin/streptomycin (pen/strep) (Gibco 15140122). Cells were maintained in T75 flasks at 37°C in a humidified incubator with 5% CO2. Passaging was performed once cultures reached 60-80% confluency. Cells were maintained up to passage 20. For long-term storage, cells were resuspended in freezing medium consisting of FBS with 10% dimethyl sulfoxide (DMSO) (Corning 25-950-CQC), gradually cooled using a Mr. Frosty container filled with isopropanol (IPA) at -80 °C, and then transferred to the vapor phase of a liquid nitrogen tank.

Newly thawed HEK293 cells were grown in T75 flasks (Corning CLS3290-100EA) and passaged at least three times prior to transfection to ensure recovery from freezing. Once cultures reach 75%-80% confluency, cells were split and seeded into 12-well plates (Corning CLS3336) or 6-well plates (Corning CLS3335) to achieve approximately 85% confluency the following day for optimal metabolic activity. On the day of transfection, confluency was verified under a microscope to ensure transfection efficiency. Four to six hours prior to transfection, the maintenance medium was replaced with Opti-MEM (Gibco 31985070). Transfections were performed using the Lipofectamine 3000 kit (Thermo Fisher L3000001) following the manufacturer’s protocol. Briefly, for each well of a 12-well plate, 1000 ng of plasmid DNA was mixed vigorously with 2µL of P3000 in 50 µL of Opti-MEM. The DNA mixture was then combined with 2µL of Lipofectamine 3000 reagent in 50 µL of Opti-MEM and incubated for 20 minutes at room temperature. A total of 100 µL of the transfection complex was added dropwise to each well of culture in 12-well plates. The whole plates were lightly shaken for even mixture before putting them back into the incubator. The reagent mixture was scaled proportionally for a 6-well plate format.

### Generation of stable tau-expressing cell line

The human 0N4R tau coding sequence with a N-terminal 3×Flag tag was excised from pcDNA3.1_Linear Tau 0N4R 3X Flag tag (Addgene plasmid # 194171 ; http://n2t.net/addgene:194171 ; RRID:Addgene_194171; gift from Stefan Stamm) (Welden, Margvelani et al. 2022) using AflII and XbaI restriction sites. The fragment was subcloned into Hygromycin B-resistant pcDNA3.1 backbone prepared by removing the ShhN insert at the identical restriction sites from pcDNA3.1 ShhN (Addgene plasmid # 37680 ; http://n2t.net/addgene:37680 ; RRID:Addgene_37680; gift from Philip Beachy). Forty-eight hours post-transfection, HEK293 cells were placed under selection pressure with 200 µg/mL Hygromycin B (Thermo Fisher Scientific). Stable expression of monoclonal cell lines was verified by immunoblotting with anti-FLAG and anti-Tau antibodies.

### Cycloheximide chase assay

At 18 h post-transfection, HEK293 cells were treated with 50 µg/mL cycloheximide (Thermo Fisher Scientific) to block de novo protein synthesis. At the indicated time points, cells were harvested and lysed on ice for 30 min in either Pierce RIPA buffer (Thermo Fisher Scientific) or sarkosyl lysis buffer (10 mM Tris-HCl, pH 7.4, 0.8 M NaCl, 1 mM EDTA, 10% (w/v) sucrose, 1% (w/v) sarkosyl) supplemented with Halt Protease and Phosphatase Inhibitor Cocktail (Thermo Fisher Scientific) and Benzonase Nuclease (EMD Millipore). Whole cell lysates using RIPA buffer were clarified by centrifugation at 16,000 × *g* for 10 min at 4 °C, and supernatant was collected. Whole cell lysates using sarkosyl lysis buffer was centrifuged at 100,000 × *g* for 1 hr at 4 °C and supernatant was collected (sarkosyl-soluble fraction). Pellet was resuspended in resuspension buffer (50 mM Tris-HCl, pH7.4, 150 mM NaCl) (sarkosyl-insoluble fraction). Protein concentrations were determined using a BCA Protein Assay Kit (Thermo Fisher Scientific).

### Drug treatment, immunoprecipitation and Western blot analysis

At 18 h post-transfection, HEK293 cells were treated with MG132 (10 µM; Tocris) or leupeptin + NH4Cl (50 µM and 10 mM, respectively) for 6 h. For immunoprecipitation of tau via 3xFLAG tag, 20 µg of the sarkosyl-insoluble fraction isolated as described in “***Cycloheximide chase assay****”* above was combined with 2 µg of anti-FLAG M2 antibody (Sigma Aldrich) in 1 ml ice-cold lysis buffer (50 mM Tris-HCl, pH 7.8, 0.5% NP40, 150 mM NaCl, 1 mM EDTA, and Halt protease/phosphatase inhibitor cocktail; Thermo Fisher Scientific). The mixture was incubated overnight at 4 °C. Next, 20 µL of Pierce Protein A/G Agarose beads (Thermo Fisher Scientific) was added to each reaction, and the mixture was incubated for 2h at 4 °C. Agarose beads were pelleted at 1000 × *g* for 1 min at 4 °C. The supernatant was removed, 1 ml of ice-cold lysis buffer was added, and pellet was washed by inverting tube several times. This wash step, including centrifugation, was repeated 4 times. After the final wash, pelleted beads were resuspended in 40 μl of 1×Laemmli sample buffer (Bio-Rad) and boiled for 5 min.

Samples were centrifuged to pellet the agarose beads, and 15 μl of the resulting supernatant was resolved on 10% PROTEAN TGX Precast Gels (Bio-Rad). Proteins were blotted onto PVDF membranes (0.2 µm pore size; Bio-Rad). Membranes were blocked with 5% (w/v) non-fat dry milk (Santa Cruz Biotechnology) in TBS containing 0.1% Tween-20 (TBS-T) for 1 h at room temperature. Membranes were subsequently probed with anti-FLAG (Cell Signaling) or anti-SUMO2/3 (Cell Signaling) antibodies overnight at 4 °C, followed by incubation with anti-rabbit HRP-conjugated secondary antibodies (1:20,000; Vector Labs) for 1 h at room temperature. Signals were visualized using Pierce ECL Western Blotting Substrate or SuperSignal West Femto Maximum Sensitivity Substrate (Thermo Fisher Scientific) on a ChemiDoc imaging system (Bio-Rad). To verify IP efficiency, membranes were stripped in Restore PLUS Western Blot Stripping buffer with rocking for 5 min at room temperature, blocked, and probed directly with HRP-conjugated anti-Mouse IgG Light Chain specific antibody (Cell Signaling) overnight at 4 °C, followed by ECL signal detection. Myc-tag, c-myc, LC3B, and p62 were detected in 10 µg of the corresponding sarkosyl-soluble samples (isolated simultaneously with the insoluble fractions) by Western Blot as described above, using their respective primary antibodies. Total protein levels obtained from Amido black (0.03 % w/v in 3% acetic acid)-stained blots were used as loading controls. Relative optical densities were determined using AlphaEaseFC software, version 4.0.1 (Alpha Innotech, San Jose, CA), normalized to total protein loaded.

### Co-immunoprecipitation

Co-IP was performed as previously described (Farrell, Kim et al. 2022). Briefly, HEK293 cells co-expressing 3xFLAG-0N4R tau and one of four myc-tagged Jade1S constructs—full-length (myc-Jade1S), a deletion mutant lacking both plant homeodomain (PHD) zinc fingers (myc-Jade1S-dd), a mutant lacking the N-terminal domain (myc-Jade1S-d1), or a fragment containing only the C-terminal domain (myc-Jade1S-d2) (Zhou, Wang et al. 2002, Chitalia, Foy et al. 2008)—were lysed in ice-cold lysis buffer containing 50 mM Tris-HCl, pH 7.8, 0.5% NP40, 150 mM NaCl, 1 mM EDTA, and Halt protease and phosphatase inhibitor cocktail (Thermo Fisher Scientific). Samples were incubated on ice for 30 min and centrifuged at 1000 × *g* for 10 min. The resulting supernatant was collected and incubated with anti-FLAG M2 antibody (Sigma Aldrich) overnight at 4 °C, and then with Protein A/G agarose beads for 2 h at 4 °C. Beads were subsequently washed and the immunoprecipitated proteins were eluted for Western blot analysis.

### Dot blot

HEK 293 cells stably expressing 3xFLAG-0N4R tau were transfected with control or myc-Jade1S expression vectors. After 24 hr, cells were washed once in ice-cold PBS, resuspended in ice-cold hypotonic lysis buffer (10 mM HEPES, pH 8.0, 1.5 mM MgCl2, 10 mM KCl) supplemented with Halt Protease and Phosphatase Inhibitor Cocktail (Thermo Fisher Scientific), and incubated on ice for 30 min. Cells were mechanically disrupted by pipetting 50 times through an extended-length, low-retention 10 µL pipette tip (Thermo Fisher Scientific). Lysates were centrifuged at 1,000 × *g* for 10 min at 4 °C, and supernatants were collected. Total protein concentration was determined by BCA assay (Thermo Fisher Scientific) and equalized across all samples.

PVDF membranes (0.2 µm pore size; Bio-Rad) were activated in 100% methanol for 1 min, rinsed in Milli-Q water for 5 min, equilibrated in TBS (50 mM Tris-HCl, pH 7.4, 150 mM NaCl) for 10 min, and placed onto TBS-wetted filter paper. Equal volumes of sample (2 µL, corresponding to 3.2 µg total protein) were spotted onto the membrane under ambient conditions. After complete absorption, membranes were blocked with 5% (w/v) non-fat dry milk (Santa Cruz Biotechnology) in TBS containing 0.1% Tween-20 (TBS-T) for 1 h at room temperature. Membranes were subsequently probed with primary conformational antibodies (anti-tau oligomer T22 or TOC1) overnight at 4 °C, followed by incubation with anti-rabbit HRP-conjugated secondary antibodies (1:20,000; Vector Labs) for 1 h at room temperature. Signals were visualized using Pierce ECL Western Blotting Substrate (Thermo Fisher Scientific) on a ChemiDoc imaging system (Bio-Rad). Membranes were stripped in Restore PLUS Western Blot Stripping buffer with rocking for 5 min at room temperature, blocked, and probed with anti-c-myc antibody (Invitrogen) overnight at 4 °C, followed by incubation with anti-mouse HRP-conjugated secondary antibodies (1:20,000; Vector Labs) for 1 h at room temperature and ECL signal detection.

### In vitro induction of tau oligomerization

*In vitro* oligomerization reactions were prepared in a final volume of 60 µL in Protein LoBind tubes (Eppendorf). Recombinant full-length human 0N4R tau (350 nM) (rPeptide) was incubated alone or with Jade1S (350 nM) in PBS (Sigma-Aldrich) supplemented with Halt Protease Inhibitor Cocktail (Thermo Fisher Scientific) for 30 min at room temperature. Tau assembly was induced by adding arachidonic acid (final concentration 2 µM; MP Biomedicals) or an equivalent volume of PBS as a vehicle control. Reactions were incubated at room temperature. At the indicated time points, 12 µL aliquots were collected into Protein LoBind tubes and immediately flash-frozen on dry ice.

### Statistics

Statistical analyses were performed using GraphPad Prism software (GraphPad Software). Quantitative data are presented as the mean ± standard error of the mean (SEM)]. Statistical comparisons between two experimental groups were evaluated using an unpaired, two-tailed Student’s *t*-test. For the cycloheximide chase assays, protein degradation kinetics were modeled using linear regression analysis. The protein half-life (*t*_1/2_) was extrapolated directly from the linear regression curve by calculating the time required for protein levels to degrade to 50% of their initial baseline (time zero) concentration. A *p*-value of less than 0.05 was considered statistically significant.

### Antibodies

Primary antibodies used for immunoblotting (IB) were: Tau12 (mouse monoclonal, 1:2,000; Millipore, Cat# [MAB2241]), HT7 (mouse monoclonal, 1:3,000; Invitrogen, Cat# [MN1000]), T22 (rabbit polyclonal, 1:1,000; Sigma-Aldrich, Cat# [ABN454]), TOC1 (rabbit polyclonal, 1:5,000; Bio-Techne, Cat# [NBP3-20162]), c-Myc (clone 9E10, mouse monoclonal, 1:1,000; Invitrogen, Cat# MA1-980), c-Myc (rabbit polyclonal, 1:6,000; Proteintech, Cat# 10828-1-AP), Myc-Tag (clone 71D10, rabbit monoclonal, 1:1,000; Cell Signaling Technology, Cat# 2278), FLAG (rabbit polyclonal, 1:1,000; Cell Signaling Technology, Cat# 2368), SUMO2/3 (rabbit polyclonal, 1:1,000; Cell Signaling Technology, Cat# 4971), LC3B (rabbit polyclonal, 1:1,000; Cell Signaling Technology, Cat# 3868), p62 / SQSTM1 (rabbit polyclonal, 1:1,000; Cell Signaling Technology, Cat# 5114), Jade1S (rabbit polyclonal, 1:5,000; Proteintech, Cat# 28472-1-AP), Mouse IgG Light Chain specific (goat, 1:2,000; Cell Signaling Technology, Cat# 91196). For immunoprecipitation (IP), anti-FLAG (clone M2, mouse monoclonal; Sigma-Aldrich, Cat# F3165) was used.

## Results

### Jade1S binds to wild-type 0N4R tau

To understand the protein-protein interaction between Jade1S and tau, we took advantage of covalent labeling mass spectrometry (CL-MS), a technique that we have previously used to characterize structural features of proteins and their interactions (Kirsch, Blake et al. 2023). We used diethylpyrocarbonate (DEPC) as our residue specific CL reagent. DEPC modifies surface-accessible nucleophilic residues (His, Lys, Tyr, Ser, Thr, Cys, and the N-terminus), which on average make up to 30% of residues in a protein (Borotto, Zhou et al. 2015). After proteolytic digestion and tandem mass spectrometry, the mass increase of the DEPC label (+72Da), can be localized with residue-level resolution. Because a residue’s reactivity towards DEPC depends on solvent accessibility and local microenvironment, changes in reactivity between sample conditions report on structural changes that occur upon ligand or protein binding (Zhou and Vachet 2012).

To identify regions of Jade1S affected by tau binding, we labeled Jade1S with DEPC in the absence and presence of the 4R tau repeat domain (Tau-RD, also known as K18; residues 244-372). Jade1S (20 uM) was prepared by itself or with Tau-RD at a 1:1 molar ratio in 10 mM MOPS (3-(N-morpholino)propanesulfonic acid buffer) at pH 7.4. Samples were then reacted with 8-fold molar excess of DEPC for 1 min at room temperature. The reaction was quenched with 50-fold molar excess of imidazole to DEPC. Labeled samples were proteolytically digested by trypsin/Lys-C and then analyzed by LC-MS/MS. The extent of DEPC modification (% mod) for residues was calculated based on the peak areas of the labeled and unlabeled peptides containing the residue of interest (Eq. 1).

Comparing these two conditions, a number of Jade1S residues showed significant changes in % mod in the presence of Tau-RD (Fig. 1A; green lines). These changes occurred in the N-terminal region of Jade1S, as well as in the PHD domains.

**Figure 1.**
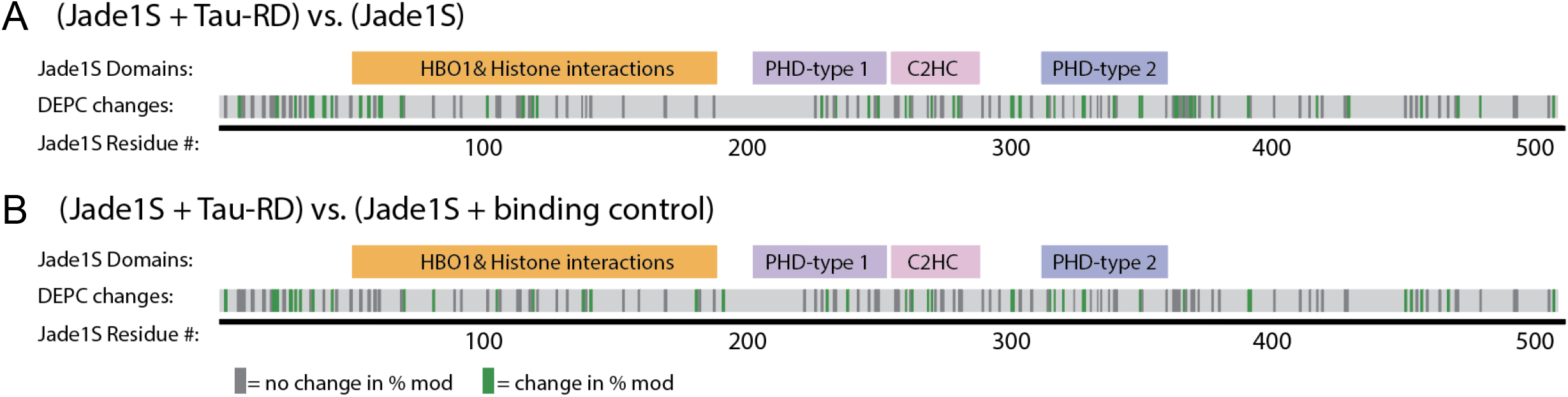
DEPC changes along the Jade1S sequence upon complex formation with Tau-RD. *A*, Jade1S % mod changes upon complex formation with Tau-RD. *B*, Changes in Jade1S DEPC modification comparing Jade1s bound to Tau-RD with Jade1S in solution with denatured cytochrome C (binding control). Green bars represent residues that are changing significantly in DEPC % mod, determined by unpaired t-test with Welch’s correction at 90% confidence interval. Dark grey bars correspond to residues that are modified by DEPC but do not change in modification extent (*n* = 2 independent experiments).

To account for changes in Jade1S labeling that do not arise from specific binding, such as DEPC consumption by the added protein (Tremblay, Kirsch et al. 2022), we also labeled Jade1S in the presence of denatured cytochrome C (CytC). CytC was chosen as a binding control due to its similarity in DEPC-reactive residues to Tau-RD, and as such would compete for DEPC to a similar extent. We then identified changes in DEPC % mod changes on Jade1S with CytC vs. Jade1S bound with Tau-RD (Fig. 1B; green lines).

Comparing these different conditions allowed us to further hone specific residues of interest that are changing due to Tau-RD complex formation. The most consistent changes were observed in the N-terminal region of Jade1S, where several neighboring residues displayed coordinated changes. Changes were also detected across the PHD domains, but these were more dispersed along the sequence. Because clustered changes in modification can reveal potential binding sites (Pianpaktr, Ramanandana et al. 2026), these data suggest that the N-terminal region of Jade1S may contribute to the interaction with Tau-RD, and the more distributed changes in the PHD domains could reflect additional contacts or conformational changes in Jade1S upon tau binding.

Next, co-immunoprecipitation was performed using HEK293 cells co-expressing 3xFLAG-0N4R tau and one of four myc-tagged Jade1S constructs: full-length (myc-Jade1S), a deletion mutant lacking both plant homeodomain (PHD) zinc fingers (myc-Jade1S-dd), a mutant lacking the N-terminal domain (myc-Jade1S-d1), or a fragment containing only the C-terminal domain (myc-Jade1S-d2). Following FLAG immunoprecipitation, we observed that 3xFLAG-0N4R tau successfully co-immunoprecipitated with full-length myc-Jade1S, myc-Jade1S-dd, and myc-Jade1S-d1, but failed to interact with myc-Jade1S-d2 (Fig. 2). Because tau retained binding to Jade1S constructs lacking either the PHD domains or the N-terminus individually, these data indicate that Jade1S likely possesses multiple binding sites for 0N4R tau, specifically within both the N-terminal and PHD domains.

**Figure 2.**
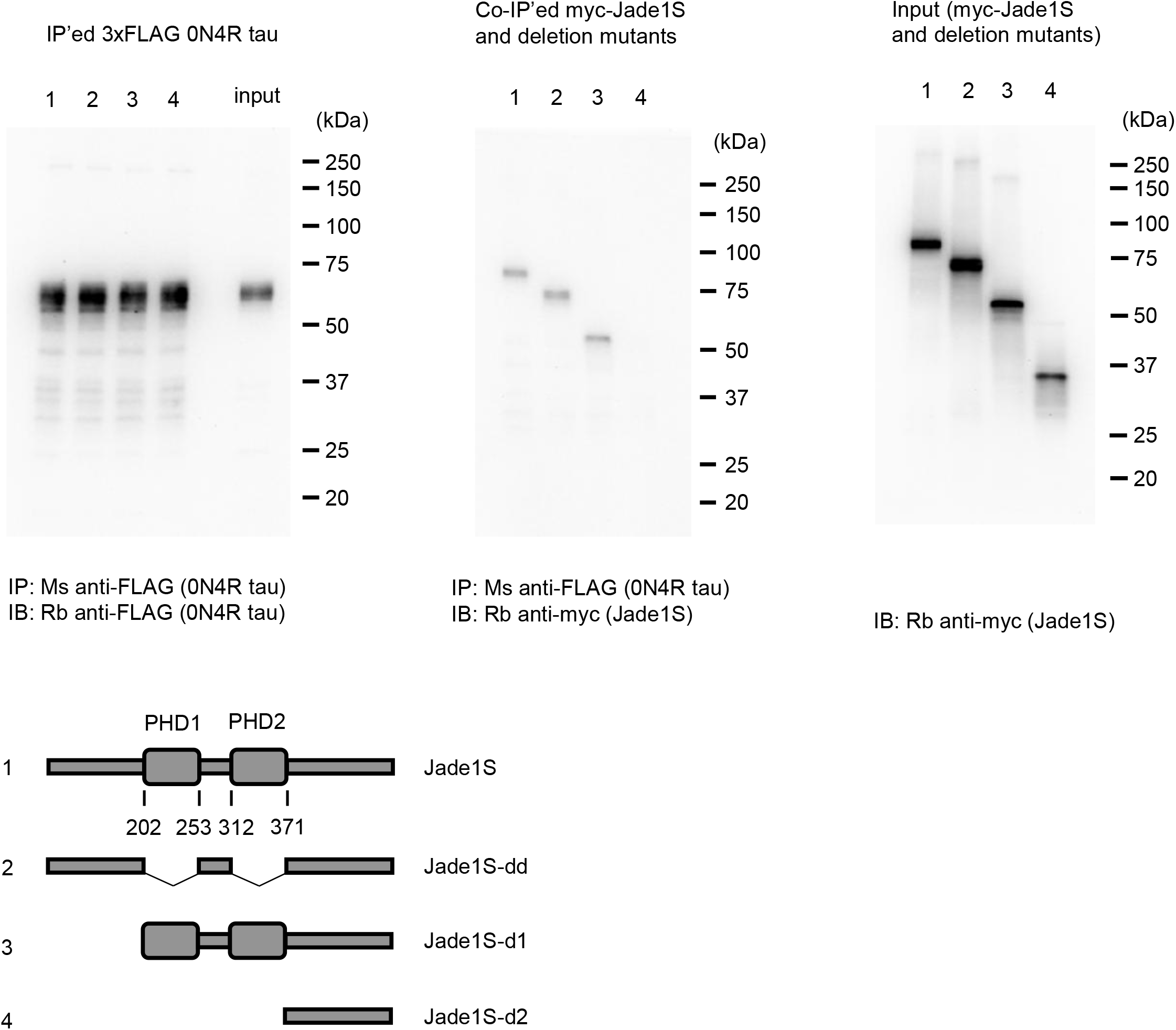
0N4R tau binds to the N-terminal and PHD domains of Jade1S. *A*, Co-immunoprecipitation (co-IP) mapping of the Jade1S and tau interaction. HEK293 cells were co-transfected with 3xFLAG-0N4R tau and one of four myc-tagged Jade1S constructs: full-length (Jade1S; lane 1), a deletion mutant lacking both plant homeodomain (PHD) zinc fingers (Jade1S-dd; lane 2), a mutant lacking the N-terminal domain (Jade1S-d1; lane 3), or a fragment containing only the C-terminal domain (Jade1S-d2; lane 4). Cell lysates (Input) were subjected to immunoprecipitation using an anti-FLAG antibody (IP: FLAG), followed by immunoblotting (IB) for myc and FLAG. 3xFLAG-0N4R tau successfully pulls down full-length myc-Jade1S, as well as the -dd and -d1 deletion mutants, but fails to interact with the -d2 fragment. This indicates that Jade1S possesses multiple, redundant binding sites for 0N4R tau located within both the N-terminal and PHD domains.

### Jade1S facilitates SUMOylation of tau

To determine whether Jade1S functions as a SUMO E3 ligase for tau, we performed *in vitro* SUMOylation assays using recombinant wild-type 0N4R tau and SUMO2. Jade1S facilitated the dose-dependent conjugation of SUMO2 to tau, peaking at a 5:1 Jade1S:tau molar ratio (500 nM : 100 nM) and yielding both mono- and di-SUMOylated tau species (Figs. 3A and 3B). In contrast, a Jade1S mutant lacking both plant homeodomain (PHD) zinc finger domains (Jade1S-dd) failed to SUMOylate tau (Figs. 3B and 3C), demonstrating that these domains are required for its catalytic activity. Assays performed with SUMO1 yielded comparable results (Figs. 3D and 3E), indicating that Jade1S can mediate both SUMO1 and SUMO2 conjugation to 0N4R tau.

**Figure 3.**
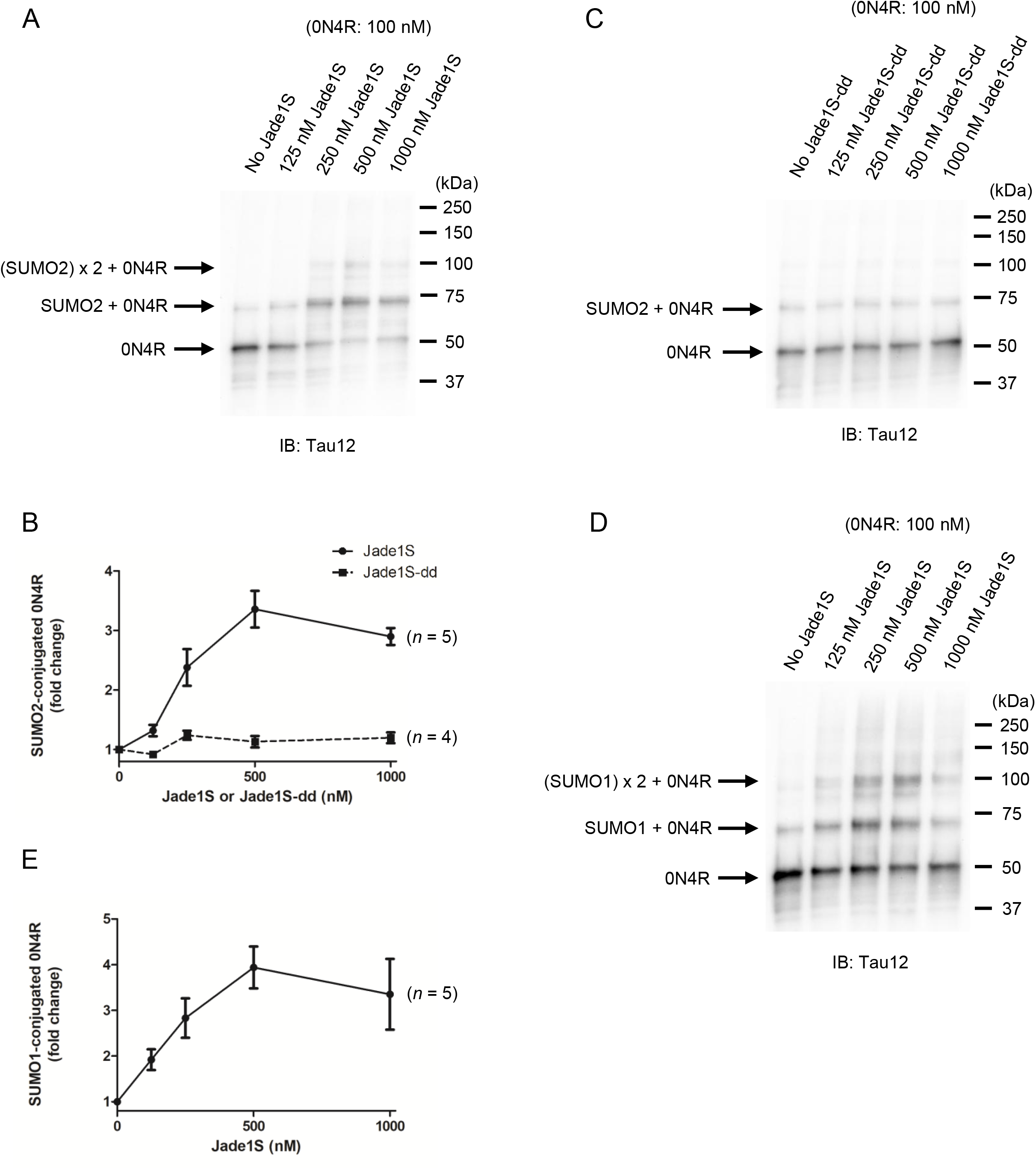
Jade1S regulates SUMOylation of wild-type 0N4R tau. *A*, Western blot analysis of an *in vitro* SUMOylation assay using recombinant SUMO2, 0N4R tau (100 nM), and Jade1S (0, 125, 250, 500, 1000 nM). De novo mono- and di-SUMOylated tau were observed. *B*, Western blot analysis of an *in vitro* SUMOylation assay using recombinant SUMO2, 0N4R tau (100 nM), and Jade1S-dd (0, 125, 250, 500, 1000 nM). Jade1S-dd lacks both plant homeodomain (PHD) zinc finger domains. No increase in SUMOylation was observed. *C*, Quantitative analysis of *A* and *B*. Jade1S facilitated the conjugation of SUMO2 to tau in a dose-dependent manner (*n* = 5), while Jade1S-dd did not (*n* = 4). *D*, Western blot analysis of an *in vitro* SUMOylation assay using recombinant SUMO1, 0N4R tau (100 nM), Jade1S (0, 125, 250, 500, 1000 nM). De novo mono- and di-SUMOylated tau were observed. *E*, Quantitative analysis of *D*. Jade1S facilitated the conjugation of SUMO1 to tau in a dose-dependent manner (*n* = 5). Data are presented as mean ± SEM.

### Tau stabilizes Jade1S

The basal half-life of Jade1S in HEK293 cells was previously reported to be 39 ± 3 min, extending to 106 ± 14 min upon co-expression with the tumor suppressor and stabilizer von Hippel-Lindau (VHL) (Zhou, Wang et al. 2002). To determine whether tau similarly stabilizes Jade1S, we performed cycloheximide chase assays in HEK293 cells co-expressing 0N4R tau and Jade1S. Tau expression markedly prolonged the estimated half-life of Jade1S to 16.9 h (Figs. 4A and 4B). To assess whether this stabilization depends on the conformational or aggregation state of tau, Jade1S turnover was measured separately in sarkosyl-soluble fractions (containing monomers and soluble oligomers) and sarkosyl-insoluble fractions (containing aggregated tau). In the sarkosyl-soluble fraction, the half-life of Jade1S was 7.2 h (Figs. 4C and 4D), whereas its turnover was arrested in the insoluble fraction (Figs. 4E and 4F).

**Figure 4.**
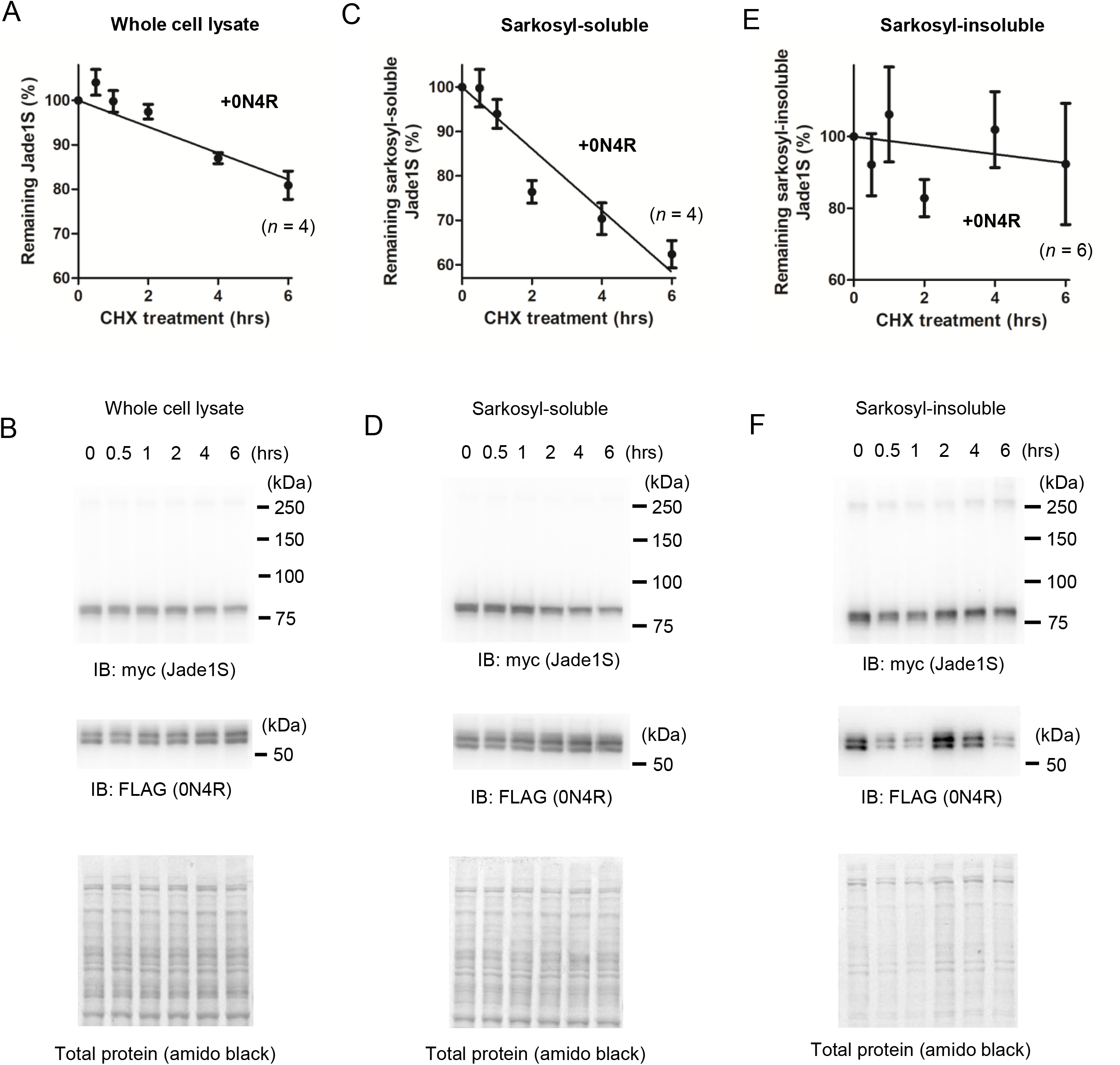
Wild-type 0N4R tau stabilizes Jade1S. *A*, HEK293 cells co-expressing 3xFLAG-0N4R tau and myc-Jade1S were treated with cycloheximide (50 µg/mL) and chased for the indicated times (0, 0.5, 1, 2, 4, and 6 h) (*n* = 4). Myc-Jade1S levels in whole cell lysate were measured by Western blot. Linear regression approximates a Jade1S protein half-life of 16.9 h when co-expressed with 0N4R tau. The basal half-life of Jade1S in HEK293 cells was previously reported to be 39 ± 3 min (Zhou, Wang et al. 2002). *B*, Representative Western blot image of myc-Jade1S from *A*. Total protein levels obtained from Amido black-stained blots were used as loading controls. *C*, HEK293 cells co-expressing 3xFLAG-0N4R tau and myc-Jade1S were treated with cycloheximide (50 µg/mL) and chased for the indicated times (0, 0.5, 1, 2, 4, and 6 h) (*n* = 4). Myc-Jade1S levels in the sarkosyl-soluble fraction were measured by Western blot. Linear regression approximates a Jade1S protein half-life of 7.2 h with 0N4R tau. *D*, Representative Western blot image of myc-Jade1S from *C*. Total protein levels obtained from Amido black-stained blots were used as loading controls. *E*, Myc-Jade1S levels in the sarkosyl-insoluble fraction were measured by Western blot (*n* = 6). Linear regression did not yield statistically significant decrease in Jade1S levels over chased time when co-expressed with 0N4R tau. *F*, Representative Western blot image of myc-Jade1S from *E*. Denaturation-resistant Jade1S-containing protein aggregates were observed (∼250 kDa). Total protein levels obtained from Amido black-stained blots were used as loading controls. Data are presented as mean ± SEM.

### Jade1S accelerates turnover of sarkosyl-insoluble tau aggregates

Tau turnover is relatively slow across biological systems, with reported half-lives ranging from ∼23 days in the human central nervous system (CNS) to ∼6.7 days in human iPSC-derived neurons and 19–32 h in HEK293 cells, depending on tags and co-factors. To evaluate whether Jade1S modulates tau stability, we performed cycloheximide chase assays in HEK293 cells expressing 0N4R tau alone or in combination with Jade1S. In sarkosyl-soluble fractions (monomers and soluble oligomers), Jade1S did not noticeably alter tau turnover rates (Figs. 5A and 5B). In contrast, Jade1S significantly accelerated the degradation of aggregated tau in sarkosyl-insoluble fractions, resulting in a ∼30% reduction relative to baseline levels over a 6-hour chase period (Figs. 5C and 5D).

**Figure 5.**
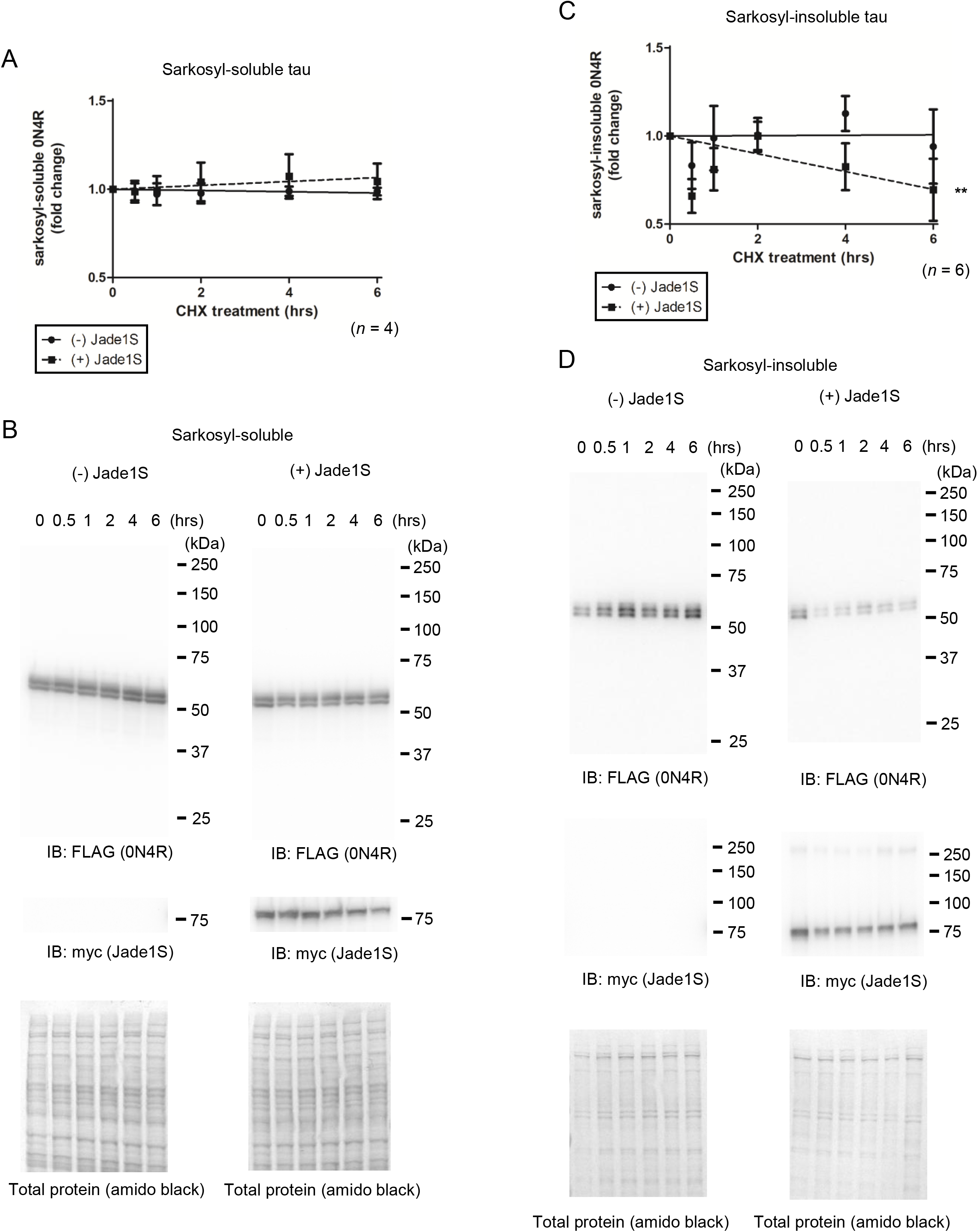
Jade1S facilitates the turnover of sarkosyl-insoluble tau aggregates. *A*, HEK293 cells expressing 3xFLAG-0N4R tau alone or in combination with myc-Jade1S were treated with cycloheximide (50 µg/mL) and chased for the indicated times (0, 0.5, 1, 2, 4, and 6 h) (*n* = 4). 3xFLAG-0N4R levels in the sarkosyl-soluble fraction were measured by Western blot. No statistically significant changes in 3xFLAG-0N4R levels were observed with or without myc-Jade1S expression. *B*, Representative Western blot images of 3xFLAG-0N4R from *A*. Total protein levels obtained from Amido black-stained blots were used as loading controls. *C*, 3xFLAG-0N4R levels in the sarkosyl-insoluble fraction from the experiments described in *A* were measured by Western blot (*n* = 6). Linear regression indicates that Jade1S co-expression led to a ∼30% reduction in sarkosyl-insoluble tau levels relative to baseline over a 6-hour chase period, while there was no statistically significant change in insoluble tau levels without Jade1S. D, Representative Western blot images of 3xFLAG-0N4R from *C*. Total protein levels obtained from Amido black-stained blots were used as loading controls. Statistical significance was assessed using linear regression to determine if the slope differed from zero. **, p < 0.01. Data are presented as mean ± SEM.

### Jade1S targets tau oligomers for degradation

To determine whether Jade1S-accelerated clearance of tau aggregates is driven by active degradation pathways, HEK cells expressing 3xFLAG-0N4R tau alone or in combination with myc-Jade1S were treated with the proteasome inhibitor MG132 or the lysosomal inhibitors leupeptin and NH4Cl. Sarkosyl-insoluble tau aggregates were immunoprecipitated via an N-terminal 3×FLAG tag and resolved by SDS-PAGE following denaturation at 95 °C in Laemmli sample buffer containing β-mercaptoethanol. This analysis revealed that the sarkosyl-insoluble fraction was composed of denaturation-resistant tau oligomers (∼400 kDa), monomers, and N-terminal fragments; notably, all species except monomers and the shortest fragments were conjugated to SUMO2/3 (Fig. 6A).

**Figure 6.**
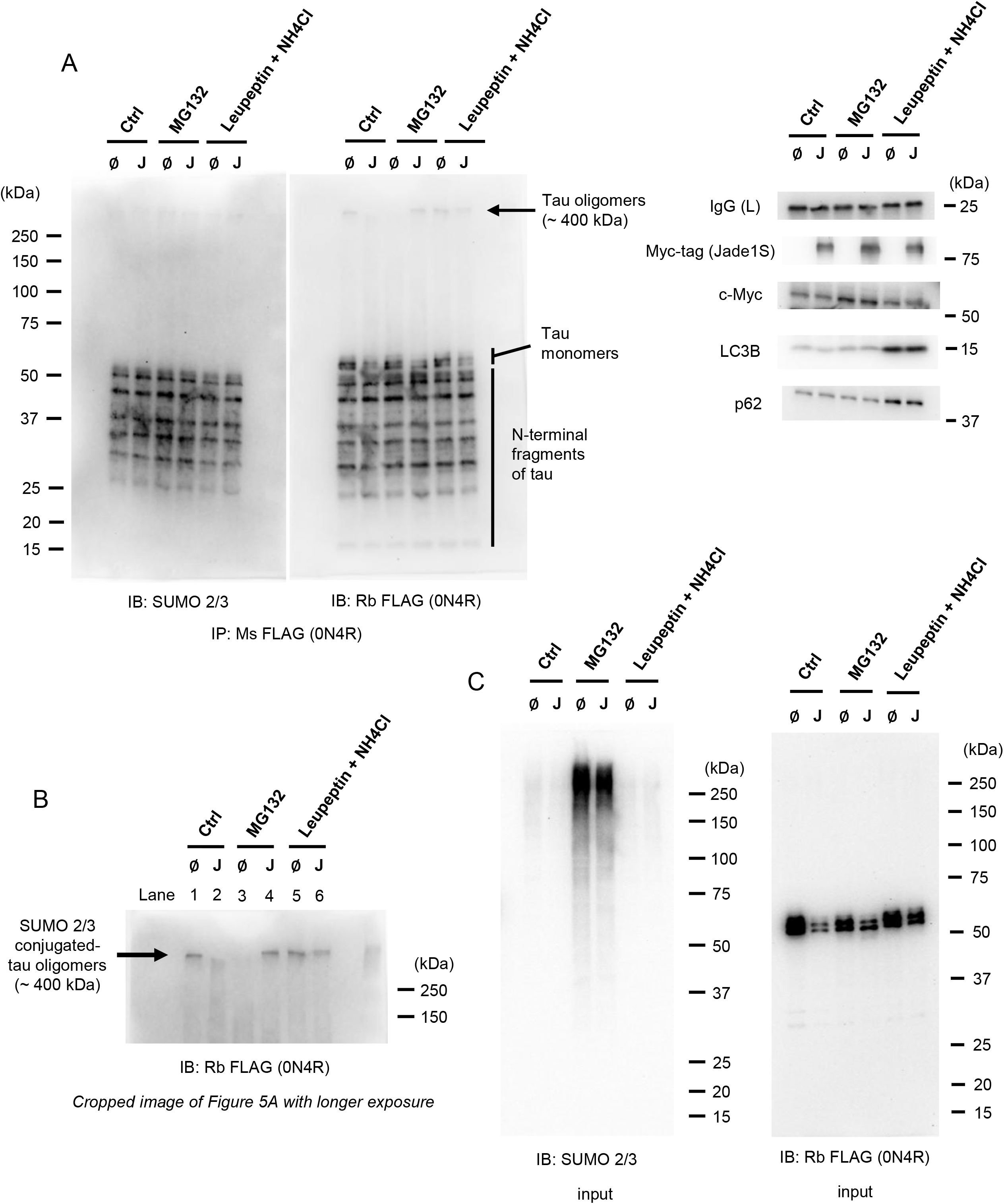
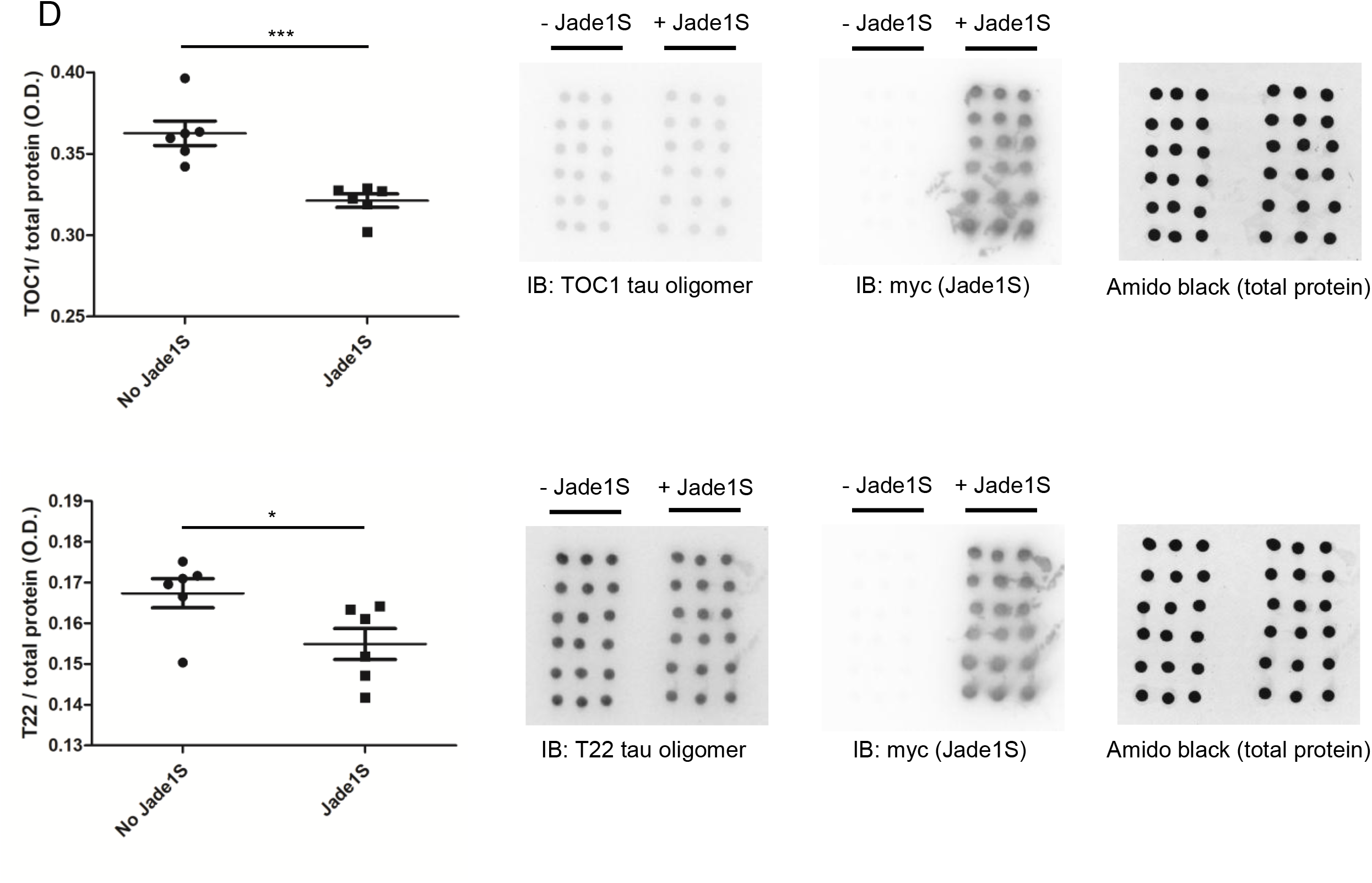
Jade1S targets tau oligomers for degradation. *A*, HEK293 cells expressing 3xFLAG-0N4R tau alone (∅) or in combination with myc-Jade1S (J) were treated with either vehicle (Ctrl), MG132 (proteasome inhibitor; 10 µM; with c-Myc as a positive control), or leupeptin + NH4Cl (lysosome inhibitors; 50 µM and 10 mM, respectively; with LC3B and p62 as positive controls) for 6 h. The sarkosyl-insoluble fraction was collected, and tau was immunoprecipitated via the 3xFLAG tag for Western blot analysis. Denaturation-resistant tau oligomers (∼400 kDa), monomers, and N-terminal fragments were immunoprecipitated as components of the sarkosyl-insoluble tau aggregates. Tau oligomers and N-terminal fragments (except the smallest one) exhibited SUMO2/3-positive signals. Myc-tag, c-Myc, LC3B and p62 were detected in the corresponding sarkosyl-soluble fraction. Mouse IgG light chain was detected by reprobing SUMO2/3-stained blot after striping. *B*, Cropped image of the anti-FLAG antibody-stained blot from *A* shown at a longer exposure. Under control conditions, SUMO2/3-conjugated tau oligomers were cleared upon Jade1S expression (lanes 1 vs. 2). Inhibition of the proteasome by MG132 rescued the oligomer band, whereas inhibition of lysosome by leupeptin + NH4Cl produced only modest recovery (lanes 2, 4, 6). In contrast, in the absence of Jade1S, baseline clearance of the tau oligomers occurred via the autophagy-lysosome pathway rather than the proteasome (lanes 1, 3, 5). *C*, Input lysate from *A* was stained for SUMO2/3 or FLAG (0N4R). Proteasome inhibition by MG132 restored broad SUMO2/3 immunoreactivity in the sarkosyl-insoluble input lysate. *D*, HEK 293 cells stably expressing 3xFLAG-0N4R tau were transfected with control or myc-Jade1S expression vectors (*n* = 6 per group). After 24 hr, cells were lysed in hypotonic buffer, and oligomeric tau levels were measured via dot blot using TOC1 and T22 antibodies. Blots show 6 independent samples per condition (vertical), performed in technical triplicate (horizontal). Total protein levels obtained from Amido black-stained blots were used as loading controls. Jade1S expression significantly lowered the levels of both TOC1- and T22-reactive tau oligomers. Statistical significance was determined using an unpaired Student’s t-test. *, p < 0.05; ***, p < 0.001. Data are presented as mean ± SEM.

Under control conditions, these SUMO2/3-modified tau oligomers were depleted upon Jade1S expression (lanes 1, 2; Fig. 6B). Proteasomal inhibition robustly rescued the oligomer band, whereas lysosomal inhibition produced only modest recovery, indicating that Jade1S targets SUMO2/3-conjugated tau oligomers for degradation predominantly through a proteasome-dependent pathway (lanes 2, 4, 6; Fig. 6B). In contrast, in the absence of Jade1S, baseline oligomer clearance occurred via the autophagy-lysosome system rather than the proteasome (lanes 1, 3, 5; Fig. 6B). Furthermore, proteasome inhibition restored broad SUMO2/3 immunoreactivity in input lysates, demonstrating that sarkosyl-insoluble aggregates are enriched in SUMO2/3-conjugated proteins destined for proteasomal clearance (Fig. 6C).

To further confirm this finding, HEK293 cells stably expressing 3xFLAG-0N4R tau were transfected with control or myc-Jade1S expression vectors. After 24 hr, cells were lysed in hypotonic buffer, and tau oligomer levels were measured via dot blot using TOC1 and T22 antibodies. Jade1S-expressing cells exhibited significantly lower levels of both TOC1- and T22-reactive tau oligomers compared to controls (Fig. 6E).

### Jade1S prevents oligomerization of tau

Because tau-induced neurotoxicity in *Drosophila* models is primarily driven by oligomerization (Wittmann, Wszolek et al. 2001, Cowan, Bossing et al. 2010, Ali, Ruan et al. 2012), and knockdown of the *Drosophila* JADE1 homolog *rhinoceros* (*rno*) exacerbated this phenotype *in vivo* (Farrell, Kim et al. 2022), we investigated whether Jade1S functions directly as a molecular chaperone to suppress tau aggregation. Exposure of recombinant 0N4R tau to arachidonic acid, an abundant brain fatty acid and established inducer of tau assembly, led to a robust increase in TOC1-immunoreactive tau oligomers *in vitro* (Figs. 7A and 7B). Notably, pre-incubating tau with Jade1S prior to the addition of arachidonic acid completely abolished oligomer formation, demonstrating that Jade1S exhibits intrinsic chaperone activity that prevents tau oligomerization (Figs. 7B and 7C).

**Figure 7.**
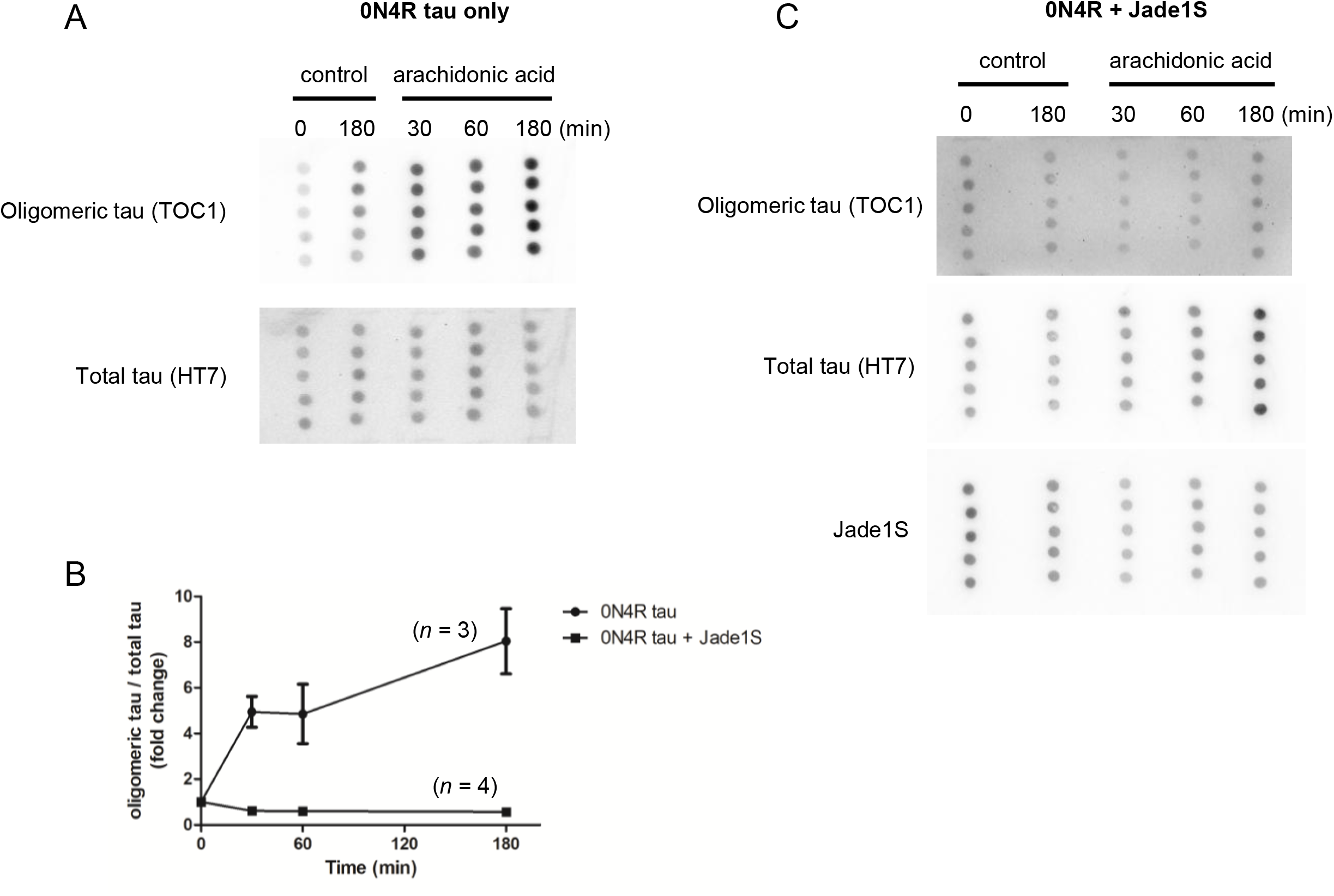
Jade1S suppresses tau oligomerization. *A*, Recombinant 0N4R tau (350 nM), alone or preincubated with Jade1S (350 nM) at room temperature for 30 min in PBS, was treated with either vehicle or arachidonic acid (tau oligomerization inducer; 2 µM) *in vitro*. Levels of TOC1-reactive tau oligomers were measured by dot blot. Representative blots show one independent sample per condition (horizontal), performed in technical quintuplicate (vertical). Total tau levels (HT7) were used as a loading control. *B*, Quantitative analysis of *A*. Arachidonic acid treatment triggered the oligomerization of tau (*n* = 3), while preincubation with Jade1S prevented this reaction (*n* = 4). Data are presented as mean ± SEM.

## Discussion

Increasing evidence implicates the dysregulated accumulation of tau as a primary driver of cognitive deficits in Alzheimer’s disease (AD) (Arriagada, Growdon et al. 1992, Warm 2003, Brier, Gordon et al. 2016). As an intrinsically disordered protein, tau depends heavily on molecular chaperones to maintain functional homeostasis and prevent pathological misfolding and aggregation (Jinwal, O’Leary et al. 2010, Freilich, Betegon et al. 2018, Mok, Condello et al. 2018). Consequently, identifying key regulators governing tau quality control represents a vital therapeutic objective. In the present study, we demonstrate that Jade1S functions as a novel SUMO E3 ligase for wild-type (0N4R) tau, suppresses tau oligomerization via intrinsic chaperone activity, and selectively targets SUMO2/3-conjugated tau oligomers for proteasomal degradation. Because the proteasomal clearance of protein aggregates typically necessitates prior disassembly, the observation that Jade1S expression drives the degradation of SUMO2/3-conjugated tau oligomers indicates that Jade1S may additionally act as a disaggregase. Furthermore, the prolonged half-life of Jade1S in sarkosyl-insoluble fractions suggests an enhanced binding affinity for aggregated tau species. Further investigation will be required to identify the specific post-translational modifications (PTMs) that modulate Jade1S chaperone activity toward pathologically misfolded tau.

Beyond its involvement in protein degradation, the broader physiological roles of tau SUMOylation remain incompletely understood. Human tau (2N4R) contains 44 lysine residues that serve as potential sites for post-translational modifications, including ubiquitination and acetylation; however, K340 (corresponding to K282 in the 0N4R isoform) remains the only established site-specific target for direct SUMO conjugation (Dorval and Fraser 2006, Luo, Xia et al. 2014, Wada, Maruyama et al. 2024). To date, only three SUMO E3 ligases for tau have been identified: PIASxα and PIASxβ, which specifically regulate conjugation of SUMO1, and TRIM11, which mediates both SUMO1 and SUMO2 modification (Zhang, Harischandra et al. 2023, Wada, Maruyama et al. 2024). Tau SUMOylation is dynamic and reversibly regulated by Sentrin/SUMO-specific proteases (SENPs) (Wada, Maruyama et al. 2024, Shui, Zheng et al. 2025). Dysregulated SUMO1 conjugation promotes tau accumulation, as SUMO1 occupancy at key lysine residues (e.g., K340) competitively blocks ubiquitination, thereby driving hyperphosphorylation, reducing solubility, and accelerating oligomeric self-assembly (Dorval and Fraser 2006, Luo, Xia et al. 2014, Takamura, Nakayama et al. 2022). Conversely, SUMO2/3 modification appears non-pathogenic; in fact, SUMO2 overexpression and its brain-penetrant mimetics have been shown to rescue long-term potentiation (LTP) and memory deficits in mouse models of tauopathy (Orsini, Pascente et al. 2024, Fioriti, Wijesekara et al. 2025). Poly-SUMO2/3 chains directly recruit the degradation machinery (Tatham, Geoffroy et al. 2008). Consistent with this mechanism, we observed enriched SUMO2/3-conjugated proteins within sarkosyl-insoluble aggregates, including tau oligomers targeted for proteasomal clearance. Notably, N-terminally SUMO2/3-conjugated tau fragments appeared to be constitutively expressed and resistant to degradation. Whether this fragment population represents a physiological feature in human neurons or an artifact of heterologous tau overexpression in HEK cells warrants further investigation.

